# RubyACRs Enable Red-Shifted Optogenetic Inhibition in Freely Behaving Drosophila

**DOI:** 10.1101/2025.06.13.659144

**Authors:** Daniel Bushey, Hiroshi Shiozaki, Yichun Shuai, Jihong Zheng, Vivek Jayaraman, Jeremy P. Hasseman, Ilya Kolb, GENIE Project Team, Glenn C. Turner

## Abstract

Optogenetic activators with red-shifted excitation spectra, such as Chrimson, have significantly advanced Drosophila neuroscience. However, until recently, available optogenetic inhibitors required shorter activation wavelengths, which don’t penetrate tissue as effectively and are stronger visual stimuli to the animal, potentially confounding behavioral results. Here, we assess the efficacy of two newly identified anion-conducting channelrhodopsins with spectral sensitivities similar to Chrimson: A1ACR and HfACR (RubyACRs). Electrophysiology and functional imaging confirmed that RubyACRs effectively hyperpolarize neurons, with stronger and faster effects than the widely used inhibitor GtACR1. Activation of RubyACRs led to circuit-specific behavioral changes in three different neuronal groups. In glutamatergic motor neurons, activating RubyACRs suppressed adult locomotor activity. In PPL1-γ1pedc dopaminergic neurons, pairing odors with RubyACR activation during learning produced odor responses consistent with synaptic silencing. Finally, activation of RubyACRs in the pIP10 neuron suppressed pulse song during courtship. Together, these results demonstrate that RubyACRs are effective and reliable tools for neuronal inhibition in Drosophila, expanding the optogenetic toolkit for circuit dissection in freely behaving animals.

## Introduction

One of the reasons *D. melanogaster* has become a powerful model for systems neuroscience is that it offers a potent pairing of optogenetic tools with genetic drivers for cell-specific expression. Optogenetics not only provides precise temporal control of neural activity but is also easier to implement and less behaviorally disruptive than other approaches, including temperature-based manipulations using *shibire* and *dTrpa1* (Bernstein et al., 2012; Hamada et al., 2008; Kasuya et al., 2009).

CsChrimson is the primary optogenetic activator used in *Drosophila* (Klapoetke et al., 2014). It is widely utilized because it can be activated by long-wavelength light (>590 nm), which can reach neurons deeper within the brain than shorter green and blue wavelengths that are more susceptible to absorption and scattering (Inagaki et al., 2014b). Additionally, because *Drosophila* photoreceptors have low sensitivity to wavelengths above 600 nm (Sharkey et al., 2020), stimulation at these longer wavelengths is less likely to interfere with behavior compared to shorter, green or blue wavelengths (Klapoetke et al., 2014).

The most commonly used optogenetic inhibitors in *Drosophila* are GtACR1 and GtACR2, both anion channelrhodopsins (ACRs) that induce membrane hyperpolarization via chloride influx. The more commonly used GtACR1 is maximally responsive to green light (515 nm), while GtACR2 is most effectively activated by blue light (470 nm) (Mohammad et al., 2017). These inhibitors have been used extensively in *Drosophila*, and—at longer wavelengths—can even be combined with visual stimulation (Mauss et al., 2017).

Red shifted ACRs have recently been isolated from Labyrinthulomycetes, a class of non-algal protists (Govorunova et al., 2020). This work identified A1ACR1 and HfACR1, two RubyACRs which exhibit peak photocurrent responses at 610 nm when fused to EYFP. Both opsins have similar properties, but HfACR1 produces stronger current. The EYFP tag is hypothesized to enhance photo-efficiency by facilitating energy transfer to the retinal chromophore, making the opsins more photo-efficient, as well as shifting the peak activation wavelength 15 nm further red.

We tested the effectiveness of these opsins in Drosophila by expressing them in neurons using both the GAL4 and LexA expression systems. We directly examined photocurrents in larval motor neurons using electrophysiological recordings, followed by functional imaging to track calcium changes in subcellular compartments of a defined neuron class in the adult visual system. Red (660 nm) light elicited strong hyperpolarizing currents with either RubyACR, but no response in GtACR1-expressing neurons. We then tested these optogenetic inhibitors in three different behavioral assays. First, we tested whether activation in glutamatergic neurons suppressed locomotion. Next, we assessed effects in dopaminergic neurons involved in associative learning. Finally we tested for optogenetic inhibition of a highly dynamic behavior, the song male flies sing to females during courtship. In all cases, RubyACR activation yielded pronounced and consistent behavioral effects consistent with neuronal inhibition.

## Results

### Functional characterization of RubyACRs in larval motor neurons

We first evaluated RubyACR performance in larval motor neurons. We used RRa-GAL4 (Fujioka et al., 2003) to drive expression of A1ACR1-EYFP, HfACR1-EYFP and GTACR1-Kir. Immunohistochemical staining revealed expression of all opsins in the motor neuron axon terminals (Figure 1A). In particular, both RubyACRs were trafficked to the plasma membrane, as their expression overlapped with a co-expressed membrane marker, myr-tdTomato (McCabe and Berthiaume, 1999). However we also observed fluorescent puncta in the cell bodies (Fig. 1 — figure supplement 1), which are likely protein aggregates that may reside within intracellular compartments. The only exception was HfACR1, where none of the three samples tested had puncta; aggregates were observed with all the other opsins tested (A1ACR: 4 out of 4 larvae containing at least one cell body with puncta; GtACR1: 1 of 2; Chrimson88: 1 of 1).

**Figure 1:**
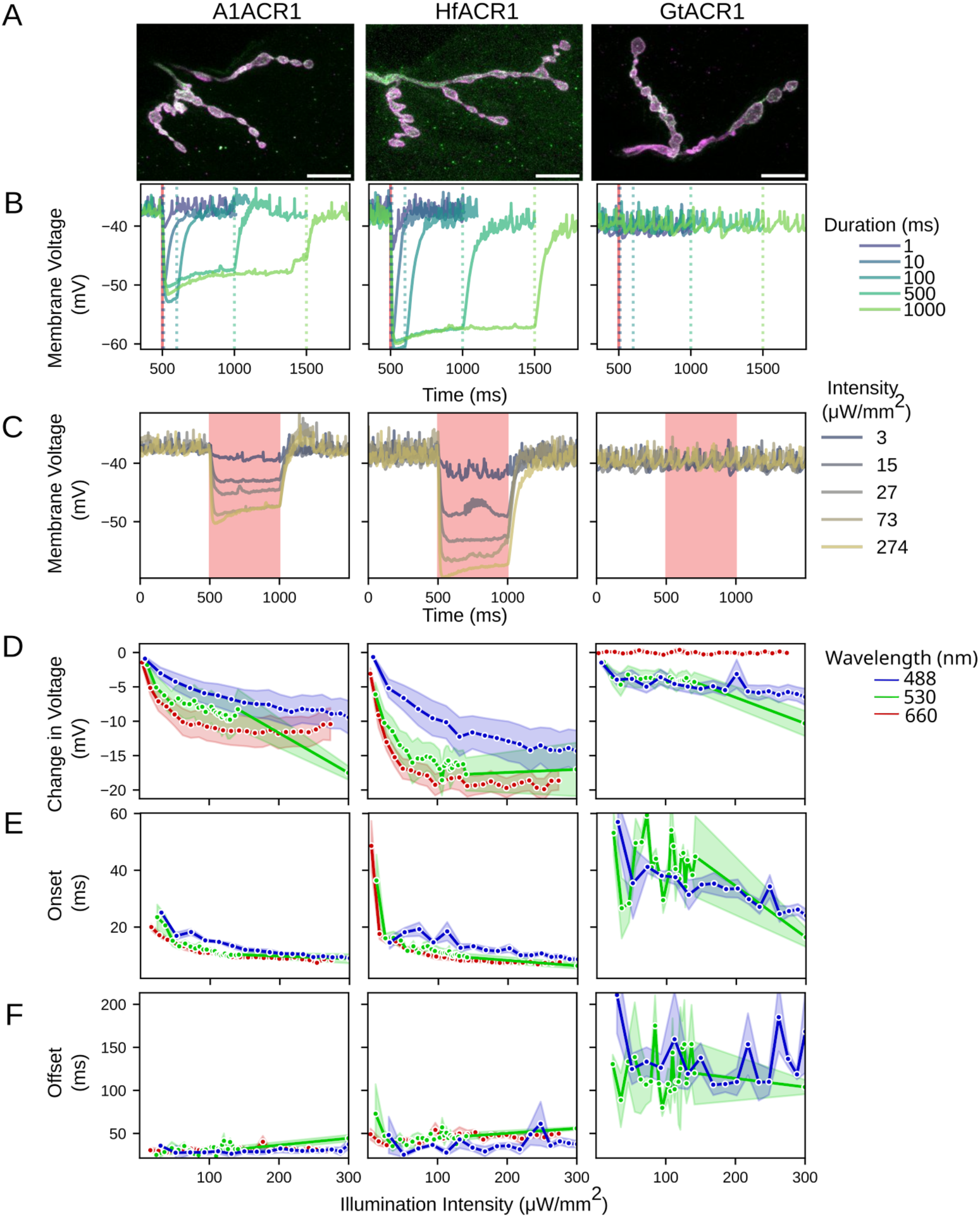
Activation of RubyACR-EYFP variants in larval motor neurons drives rapid and robust hyperpolarization. A. Expression of ACR variants at the neuromuscular junction. Green shows staining for the ACR, magenta is the membrane localized myr::tdTomato and white represents colocalization; all images are maximum intensity projections (scale bar = 10 μm). Columns correspond to ACR variants: A1ACR1 (left), HfACR1 (middle), and GtACR1 (right). This organization is maintained across panels B–F. B. Membrane potential responses to 660 nm light (270 μW/mm²) at varying pulse durations (1, 10, 100, 500, and 1000 ms, as indicated in the legend). Red vertical line marks illumination onset; colored dotted lines represent illumination offset for each stimulus duration. C. Mean membrane voltage responses to 500 msec stimulation across 25 intensities ranging from 4 to 274 μW/mm² in 11 μW/mm² increments. D. Mean change in membrane potential in response to stimulation at different wavelengths (488 nm blue, 530 nm green, and 660 nm red) and intensities. Line color indicates wavelength; shading represents standard error. E. Mean half onset time, defined as the time (ms) at which the membrane potential crosses the halfway point between pre-illumination baseline and the minimum voltage during illumination. F. Mean half offset time, defined as the time (ms) at which the membrane potential crosses the halfway point between the voltage at illumination offset and the return to baseline voltage. Sample sizes through A-F: A1ACR1-EYFP: n = 10 recordings; HfACR1-EYFP: n = 10; GtACR1-kir: n = 11.

We characterized the electrophysiological responses of motor neuron cell bodies expressing different inhibitory opsins. Opsin activation was driven by a 660 nm LED; although this wavelength is on the shoulder of the activation spectrum of the RubyACRs (peak activation at 610 nm), it is also near the limits of the visual acuity of Drosophila (Sharkey et al., 2020), which should minimize behavioral effects of the light alone. Testing light pulses of varying duration (Figure 1B), we found that pulses as brief as 10 msec elicited reliable, temporally precise hyperpolarization with both RubyACRs but not with GtACR1. Consistent with previous reports, HfACR1 exhibited larger hyperpolarization amplitudes than A1ACR1 (Govorunova et al., 2020). Longer pulse durations produced sustained hyperpolarization with minimal adaptation over the tested timescales. In contrast, GtACR1 did not elicit detectable hyperpolarization at 660 nm for any tested pulse duration.

We next evaluated responses to varying red light intensities. With 660 nm illumination, both RubyACRs elicited a clearly detectable response to a 500 msec pulse at just 4 μW/mm² (Kruskal– Wallis test *p* < 0.05; Wilcoxon U test: *p* = 0.008 for A1ACR1, *p* = 0.002 for HfACR1), and responses approached saturation at 62 μW/mm² (Figure 1C,D). Even at the highest tested intensity (274 μW/mm²), GtACR1-expressing neurons showed no response, as expected from its spectral sensitivity.

RubyACRs tagged with EYFP have a broad action spectrum in vitro (Govorunova et al., 2020). We observed a correspondingly broad sensitivity in vivo, where illuminating with blue (488 nm) and green (530 nm) LEDs triggered hyperpolarization for both opsins (Figure 1D). Nevertheless, red illumination tended to produce a greater hyperpolarization than the other wavelengths. By contrast, although red light did not elicit a response with GtACR1, both blue and green light did trigger hyperpolarization. Compared to GtACR1, the RubyACRs produced a greater hyperpolarization across all the tested wavelengths, including at their respective optimal wavelengths. With green illumination, GtACR drove a maximum decrease in mean membrane potential of 13 mV, whereas the maximum change for A1ACR1 and HfACR1 with red light was 18 mV and 21 mV respectively.

RubyACRs exhibited faster onset activation and offset deactivation kinetics in response to blue and green illumination compared to GtACR1 (Figure 1E,F). For statistical comparison, we pooled all responses to 530 nm light at intensities above 46 μW/mm^2^. The time to half of the maximum hyperpolarization was 8.4 msec for A1ACR, 8.0 msec for HfACR and 20 msec for GtACR1 (GtACR1 vs. A1ACR1 Mann-Whitney U test 2.3E-15; GtACR1 vs. HfACR1 Mann-Whitney U test 2.4E-20). The small difference between the two RubyACRs was also significant (Mann-Whitney U test p = 0.02). Offset times were also faster, with half times of 43 msec for A1ACR and 50 msec with HfACR1, while GtACR1 had an half offset of 120 msec (GtACR1 vs. A1ACR1 7.3E-48; GtACR1 vs. HfACR1 1.3E-50; A1ACR vs. HfACR1 p= 9.9E-12 Mann-Whitney U test). HfACR1 was significantly slower than A1ACR (9.9E-12 Mann-Whitney U test).

Together, these results demonstrate that RubyACRs outperform GtACR1 in terms of both the amplitude and kinetics of the evoked hyperpolarization.

### RubyACR drives hyperpolarization in ex vivo brains

In principle, red-shifted anion channelrhodopsins offer the ability to combine optogenetic manipulation with simultaneous calcium imaging using GCaMP. To test this, we targeted the fly visual system, focusing on Mi1 neurons—non-spiking interneurons involved in motion detection. Mi1 neurons extend dendrites into layers 1 and 5 of the medulla and project axons to layers 9 and 10 (Hasegawa et al., 2011).

We co-expressed either A1ACR1 or Hf1ACR with GCaMP6s in Mi1 neurons and recorded responses to 660 nm light pulses in ex vivo brains. Using two-photon imaging (920 nm excitation), we monitored GCaMP signals from regions of interest (ROIs) located on axonal projections to medulla layers 9–10 (Figure 2A). Optogenetic stimulation was delivered through the microscope’s optical path, illuminating the entire field of view continuously for 20 sec with a 660 nm LED.

**Figure 2:**
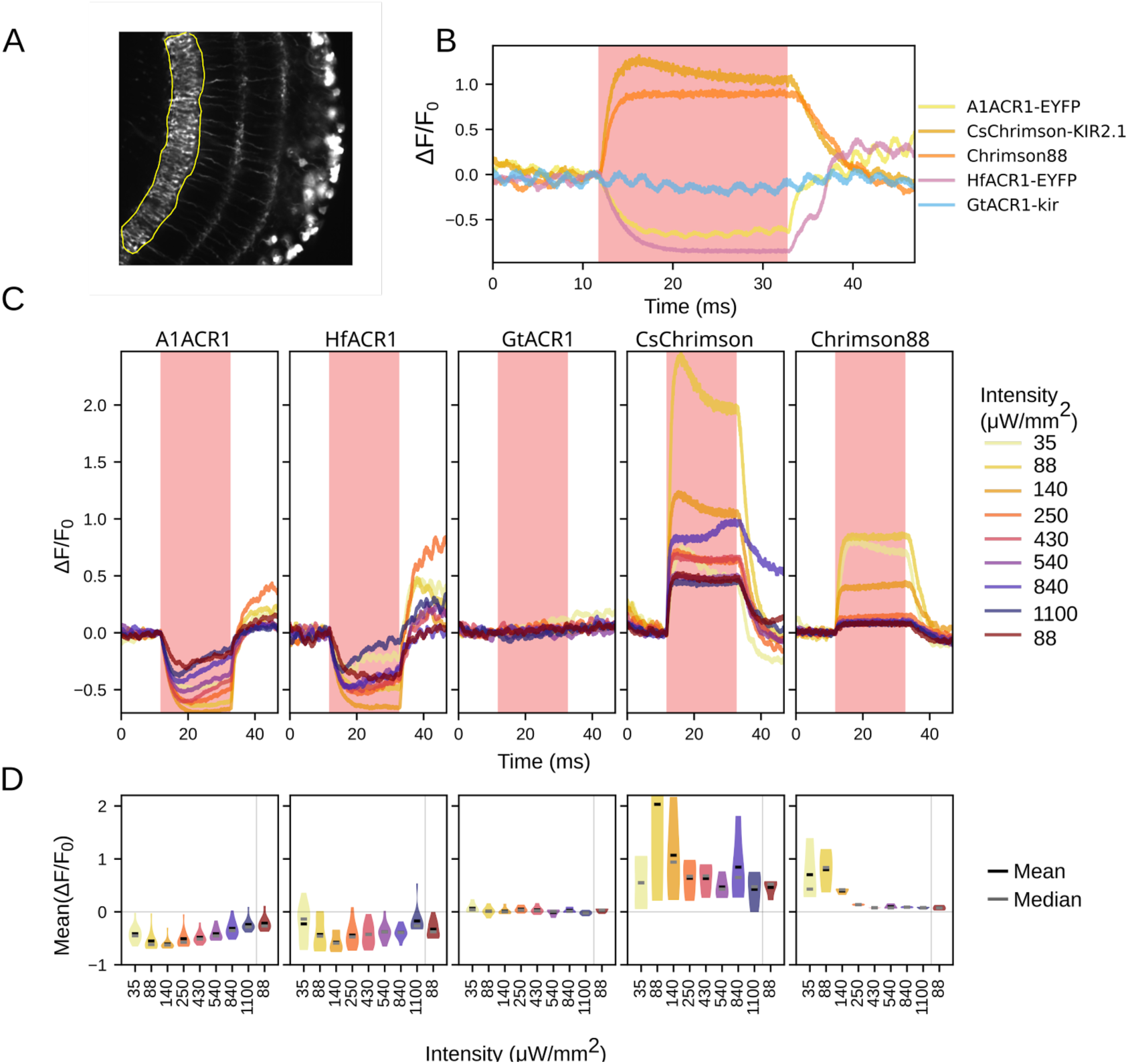
Measuring RubyACR driven inhibition in Mi1 neurons. A. Maximum intensity projection over time from two-photon imaging of Mi1 neurons expressing GCaMP6s and A1ACR1-EYFP. Fluorescence changes were measured in medulla layers 8–10, outlined in yellow. B. ΔF/F₀ traces from individual animals. The red shaded region indicates the 30-second period of 660 nm illumination at 88 μW/mm². C. The ΔF/F₀ response time course as each ChR is illuminated with increasing 660 nm light intensities over the 20 sec period (shaded in red). Legend order corresponds to testing order. Light intensity was increased sequentially across trials, except for the final trial (brown), in which intensity was returned to 88 μW/mm². D. Mean ΔF/F₀ values over the 20 sec stimulation period from the data in C. Intensity order follows the order in which the illumination intensities were applied. All responses from A1ACR1- and HfACR1-expressing flies were statistically significant (p<0.05 from Kruskal– Wallis test followed by pairwise Mann–Whitney U tests). The only exception to this was HfACR1 at 1100 μW/mm². This observation, together with the U-shape of both RubyACR curves is an indication of channel desensitization at higher intensities. This was confirmed by the smaller amplitude of the final response to the repeat of 88 μW/mm^2^ stimulation (A1ACR1 Kruskal–Wallis test *p* = 9.7E-10; Wilcoxon U test: *p* = 3.9E-3, HfACR1 Kruskal-Wallis test p = 0.02: *p* = 0.088). Sample sizes through A-D: A1ACR1-EYFP: n = 12; HfACR1-EYFP: n = 10; GtACR1-kir: n = 6; CsChrimson-KIR2.1: n = 4, and Chrimson88: n = 3.

Illumination intensity increased stepwise from 35 to 1100 μW/mm² with each trial. Changes in GCaMP6s fluorescence were detected over the constant baseline fluorescence produced from the EYFP tag. With both A1ACR1 and Hf1ACR, GCaMP fluorescence significantly decreased during 660 nm illumination (Figure 2 B,C A1ACR1 Kruskal-Wallis test p=3.1E-14, post-hoc Mann Whitney U test A1ACR1 p<3.2E-3; HfACR1 Kruskal-Wallis test p=2.9E-11, post-hoc Mann Whitney p<0.012); response amplitudes did not differ between the two RubyACRs (Kruskal– Wallis test *p*=3.4E-11; pairwise Mann–Whitney U tests p > 0.05 ). In contrast, GtACR1 did not elicit any measurable change in GCaMP signal under the same stimulation conditions (Wilcoxon signed-rank test, *p* > 0.05). As expected, when either CsChrimson or Chrimson88 were present, activation increased GCaMP fluorescence.

In RubyACR-expressing neurons, GCaMP6s signals progressively decreased as illumination intensity increased from 35 to 140 μW/mm² (Figure 2D). However, at intensities above 140 μW/mm², the magnitude of the GCaMP response declined, which is consistent with previous findings that high intensity light desensitizes these channels (Govorunova et al., 2020; Sineshchekov et al., 2023). Supporting this interpretation, repeating the 88 μW/mm² stimulus at the end of the protocol elicited a reduced response (A1ACR1 Kruskal–Wallis test *p* = 9.7E-10; Wilcoxon U test: *p* = 3.9E-3, HfACR1 Kruskal-Wallis test p = 0.02: *p* = 0.088). Desensitization was not unique to RubyACRs, as both CsChrimson and Chrimson88 also produced weaker responses when illumination intensity exceeded 140 μW/mm², demonstrating a similar profile.

Our findings demonstrate that simultaneous GCaMP imaging and optogenetic inhibition are feasible using RubyACRs. Moreover, the reduction in axonal calcium levels upon RubyACR activation is consistent with decreased neuronal excitability, which would in turn reduce synaptic release.

### Optogenetically inhibiting walking with RubyACRs

Given the effectiveness of RubyACRs in hyperpolarizing motor neurons in the larvae, we next tested whether they would inhibit motor behavior in freely behaving adult flies. We quantified the strength of optogenetic inhibition by measuring walking speed in the Fly Bowl assay (Robie et al., 2024, 2017). In this setup, individual flies are tracked as they navigate a circular arena using an infrared camera system (850 nm illumination; Figure 3A). Locomotor activity was recorded while flies were exposed to increasing intensities of stimulation light delivered by LEDs positioned beneath the Fly Bowl—a favorable configuration for targeting leg motor neurons.

**Figure 3:**
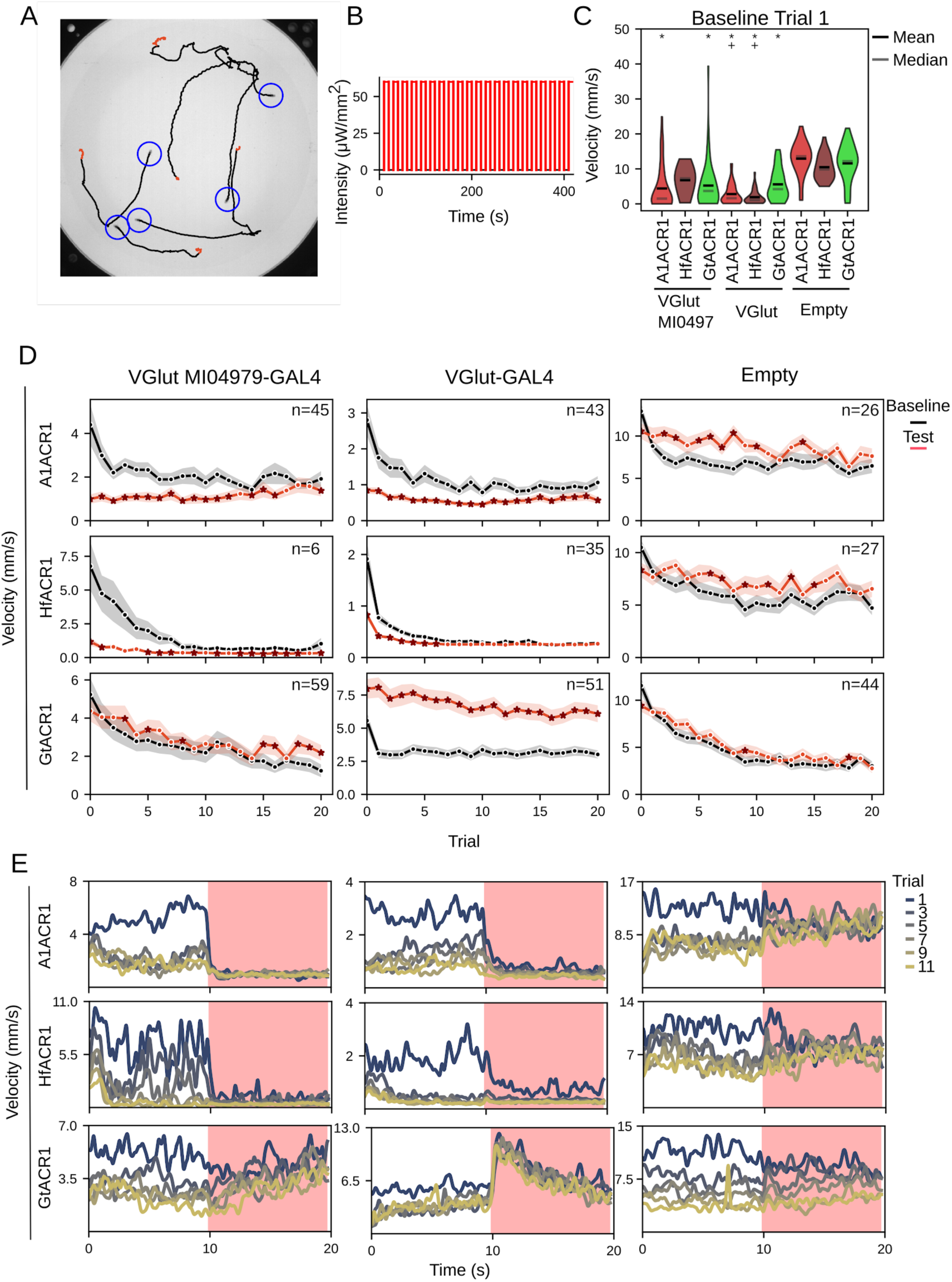
Optogenetically inhibiting spontaneous motor behavior. A. Example image from the Fly Bowl showing fly positions and their trajectories. Blue circle marks the initial position of a fly. Black traces represent movement during the 10 sec baseline period, and red traces show movement during the 10 sec testing period under 660 nm LED illumination at 29 μW/mm². Flies shown are VGlut MI04979-Gal4 driving 20XUAS-A1ACR1. B. Stimulation protocol: each trial consists of a 10-second baseline period (black) followed by a 10-second testing period (red), during which the chamber is illuminated with 660 nm light at 29 μW/mm². This cycle is repeated for 21 trials. C. Walking speed during the baseline period of the first trial, prior to any LED stimulation. Asterisks indicate driver–effector genotypes with significantly lower speed than their empty-driver controls (p < 0.05, Kruskal-Wallis test with post hoc Mann-Whitney U test). A plus symbol (+) denotes a significant difference between GtACR1 and other ChRs expressed with the same driver (p < 0.05). D. Mean velocity for each trial. Black lines show average velocity during baseline; red lines show velocity during the subsequent illuminated testing period. Each row corresponds to a specific ChR variant, and each column to a specific driver (columns continue into panel E). Asterisks as markers denote cases where Kruskal-Wallis analysis found a significant difference (p < 0.05), followed by a Wilcoxon signed-rank test comparing baseline to testing periods (p < 0.05). Numbers of flies tested are indicated in the right top corner of each plot. E. Mean velocity measured per second during trials 1, 3, 5, 7, 9, and 11, as indicated in the legend. The red shaded region indicates the illumination during the testing period. ChR variants are arranged along the rows, columns match the drivers shown in panel C.

We tested two different GAL4 lines designed to drive expression in glutamatergic neurons, thereby targeting motor neurons but also numerous other cells in the fly brain. This broad expression did result in some lethality with all three ACRs, as well as the widely used optogenetic activator Chrimson88 (Table 1; Strother et al., 2017). The survival rate was highest for GtACR1 and lowest for HfACR1. Additionally, we noticed that simply expressing the opsins decreased baseline locomotor activity compared to empty driver lines; for example, with VGlut MI04979 A1ACR was 65%, HfACR1 35% and GtACR1 54% lower than the corresponding empty driver control (Figure 3C).

**Table 1:**
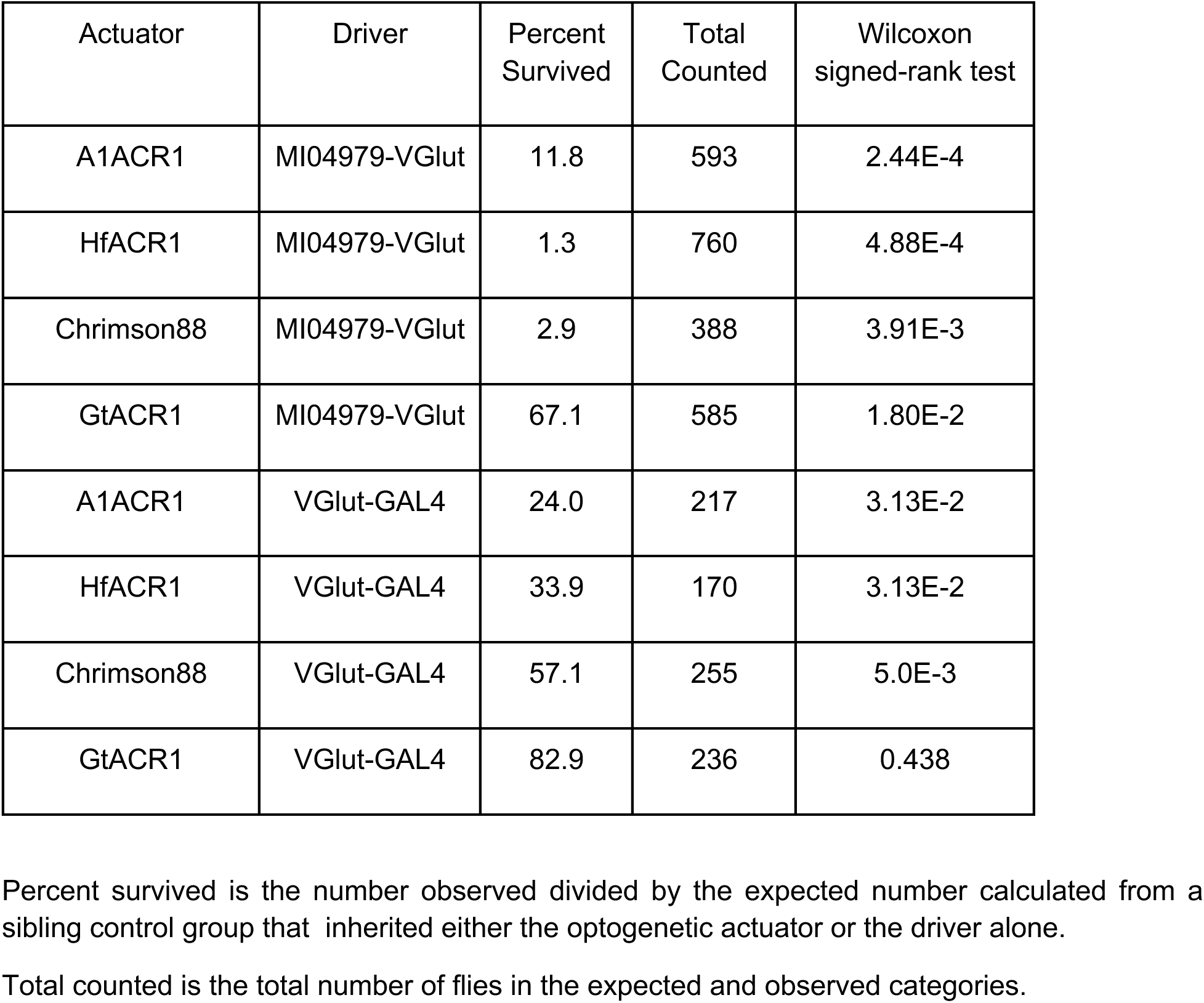
Percent Survival of Flies Expressing Different Optogenetic Actuators.

We tested the effects of red light on locomotion using a protocol that consisted of alternating 10-second baseline and 10-second illumination periods (Figure 3B). This cycle was repeated for a total of 21 trials, using 660 nm light at a constant intensity of 60 μW/mm². Red light reduced walking speed in RubyACR-expressing flies compared to both empty-driver controls and GtACR1-expressing animals. This reduction began immediately upon light onset and remained stable throughout the illumination period (Figure 3E). Although walking speed increased after the light was turned off, it did not return to pre-stimulation levels. As a result, walking speed during the ‘baseline’ periods of successive trials gradually declined. This effect was especially pronounced in flies expressing HfACR1 under VGlut-GAL4, where baseline and test-period speeds eventually became indistinguishable. We also tested LexAop2-driven RubyACR lines and observed similar results: red light suppressed locomotor activity, and this suppression persisted into the baseline period of the next trial (Fig. 3 —figure supplement 1). The rapid recovery shown in our recordings of larval motor neurons suggest that the persistent suppression does not arise in the motor neurons bearing the opsin; it may be that simply repeatedly inhibiting the fly’s locomotion circuit (or the extensive set of glutamatergic neurons these drivers target), changes the properties of the network and leads to persistent behavioral effects. Overall however, RubyACRs consistently inhibited spontaneous locomotion with a variety of drivers and expression systems.

### Evaluating RubyACR efficacy with a dopamine-dependent learning paradigm

Studies using GtACR2 in mouse cortical neurons have demonstrated that activation hyperpolarizes the soma but, unexpectedly, depolarizes presynaptic terminals (Emiliani et al., 2022; Mahn et al., 2016; Malyshev et al., 2017; Messier et al., 2018; Rost et al., 2022). This effect is attributed to an inverted chloride gradient at terminals, so light-induced channel opening actually leads to a chloride efflux, depolarizing terminals and promoting neurotransmitter release. In contrast, the reduced calcium signals we observed in the axonal arbor of Mi1 cells upon RubyACR activation indicate that presynaptic sites in Drosophila are hyperpolarized

To further validate RubyACRs as effective inhibitory tools in this system, we used a behavioral assay that provides a sensitive measure of synaptic release. Dopamine release from the PPL1-γ1pedc neurons that innervate the fly mushroom body is critical for aversive olfactory conditioning (Aso and Rubin, 2016). Pairing presentation of an odor with optogenetic activation of PPL1-y1pedc neurons produces a robust aversive association, causing flies to subsequently avoid that odor.

We expressed various opsins in PPL1-γ1pedc neurons using the MB320C split-GAL4 driver and trained flies using red light pulses at two intensities (13 or 26 μW/mm²; 625 nm peak emission) paired with odor presentation. As anticipated, CsChrimson activation induced aversive memory, and flies avoided the paired odor (Figure 4C). Activation of RubyACRs actually led to the opposite behavioral outcome, with flies exhibiting a moderate preference for the paired odor during testing. While we did not specifically predict appetitive memory formation in this neuron, this result aligns with prior studies demonstrating that suppressing/blocking synaptic release in other dopaminergic neurons during training can switch the learned association from aversive to appetitive (Yamagata et al., 2016). For comparison, flies expressing GtACR—which is not activated by the red light— displayed no odor preference after training.

**Figure 4:**
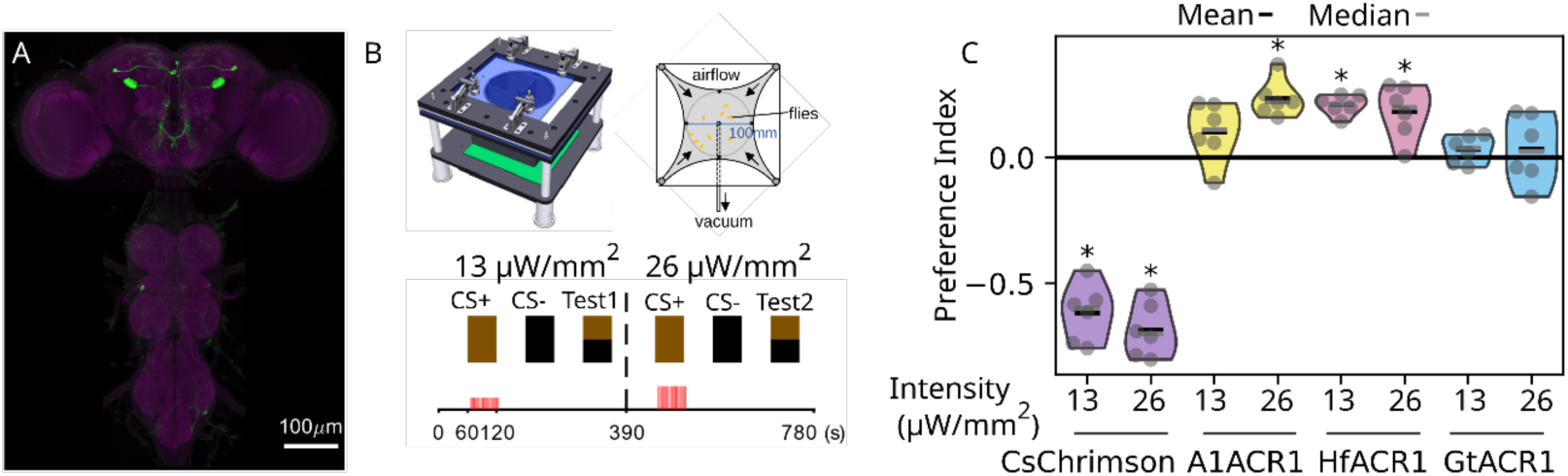
Optogenetic activation of RubyACRs and CsChrimson drives opposing outcomes in associative conditioning. A. MB320C labels punishment-signaling dopaminergic neurons innervating the γ1pedc compartment of the MB. Expression pattern of CsChrimson-mVenus (green) shown with neuropil counterstaining of Brp (magenta). B. Schematic of the optogenetic olfactory learning assay (adapted from (Aso and Rubin, 2016)). Flies underwent two rounds of odor-light pairing using 629 nm LED illumination at 13 and 26 μW/mm². Each pairing was followed by a test for preference between the LED-paired odor (conditioned stimulus, CS+) and a control odor (CS-). C. Preference index (PI) of flies expressing different optogenetic inhibitors (ACRs) or activators (CsChrimson) under MB320C. Positive PI values indicate appetitive memory; negative values indicate aversive memory. Each group includes n = 6 replicates (dots). Black bars show the mean, and gray bars indicate the median. Asterisks denote trials with a significant response relative to zero (p = 6.0E-06, Kruskal-Wallis test across groups followed by one-sample Wilcoxon signed-rank tests within groups).

Collectively, these findings validate RubyACRs as reliable optogenetic inhibitors of synaptic output in intact neural circuits.

### RubyACRs effectively inhibit courtship song circuitry

To test for optogenetic inhibition of a different, highly dynamic behavior, we examined courtship song. Males sing to females as part of the courtship behavior, with the song composed of two auditory elements: pulses which come from short, strong wing vibrations and sine song which are more prolonged continuously varying vibrations. The pIP10 neuron is a central node in the song circuit - activating this neuron strongly increases pulse song and weakly increases sines while inactivating the cell reduces both song types (Ding et al., 2019; Lillvis et al., 2024). We expressed the RubyACRs selectively in the pIP10 neuron to assess whether their activation could suppress pulse song.

Male-female pairs were placed in a chamber with a microphone to record courtship song during baseline, testing, and post periods, each spanning 5 sec (Figure 5A,B). We tested a series of red light (629 nm) stimulations of increasing intensity on pulse song frequency. Both A1ACR1 and HfACR strongly suppressed pulse song frequency in an intensity-dependent manner, with inhibition plateauing at ∼17 μW/mm² (Figure 5C). At this intensity, pulse frequency was sharply reduced during testing compared to baseline but fully recovered during the post-testing period (Figure 5D; A1ACR1 testing: -85%, post-testing: 3.4%; HfACR1 testing: -84%, post-testing: - 32%). The reductions during the testing period were statistically significant compared to empty-driver controls; this comparison was the most appropriate because higher light intensities modestly reduced pulse frequency even in control animals (Fig. 5—figure supplement 1A). In contrast, the small reduction in singing observed in GtACR1-expressing flies was not significantly different from the reduction in empty-driver controls (Figure 5D, Kruskal–Wallis test *p*=1.0E-3; pairwise one tail Mann–Whitney U tests p > 0.88 testing for decreased pulse counts in the driven line compared to controls for each illumination intensity).

**Figure 5:**
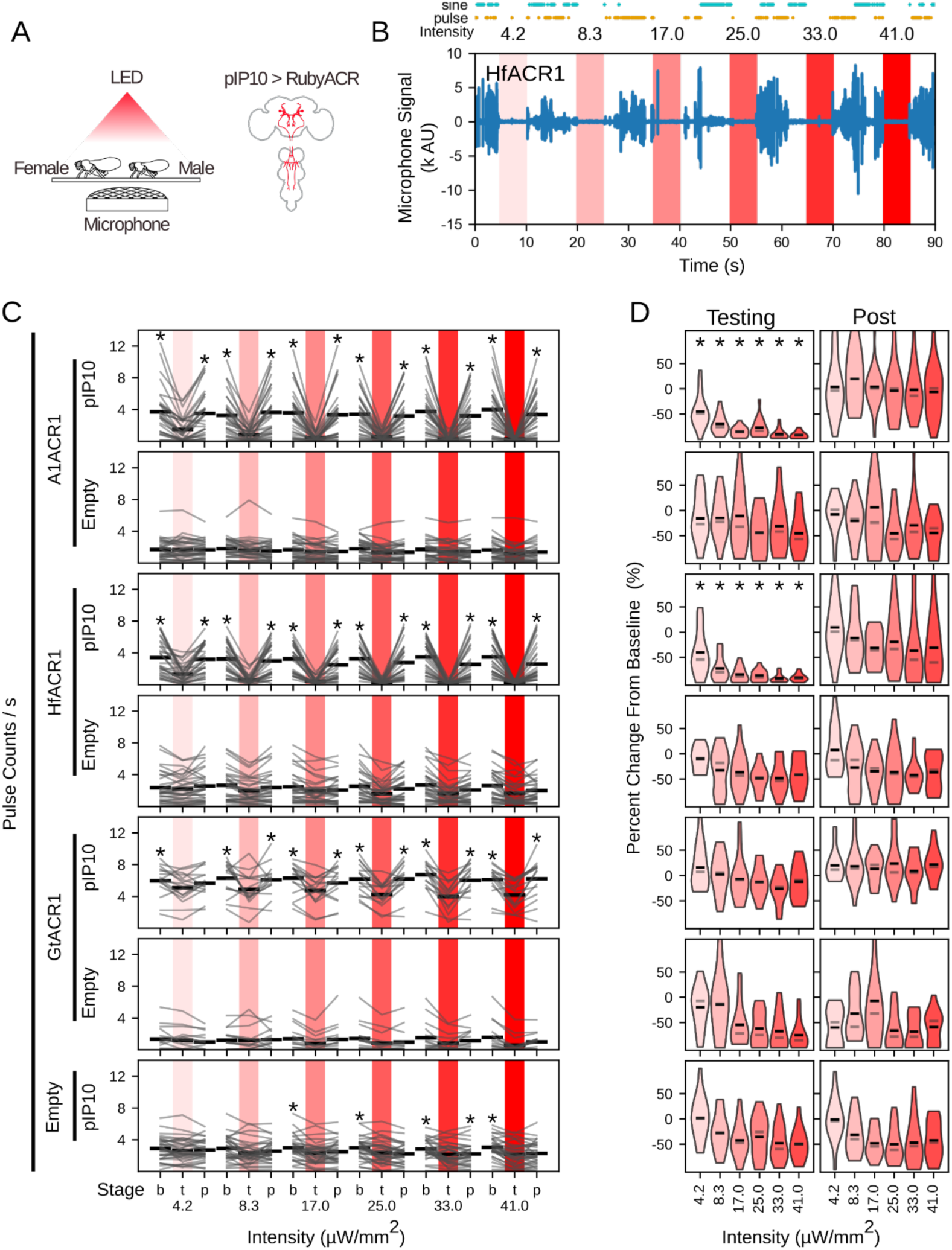
Activating RubyACRs in pIP10 neurons suppresses courtship pulse song. A. Left: Diagram of the courtship assay. Singing male flies are recorded using a microphone while the chamber is illuminated with a 629 nm LED. Right: Schematic of the pIP10 neuron within the central nervous system. B. Example audio recording from a single fly expressing HfACR1 in pIP10. Red shading indicates periods of LED illumination. Illumination intensity increases from left to right as indicated. Courtship song elements (sine and pulse) are labeled above the trace. C. Mean rate of song pulses for individual courting pairs over trial stages. Each row represents a particular driver-effector combination, as indicated at left. Stages are 5-second epochs of: baseline (b) prior to illumination; testing (t) when illuminated and post-testing (p). Illumination intensity increased with each successive trial as indicated by the shading. Gray lines represent data from individual courting pairs (n = 16), black bars indicate mean over all pairs. Asterisks denote significant differences between the testing period and either the baseline or post period (p < 0.05, Kruskal-Wallis test with post-hoc Wilcoxon signed-rank test). D. Percent change in pulse song frequency from baseline to testing (left) and from baseline to post (right), based on data in **C**. Driver-effector combinations for each row continue from **C**. Violin plots reflect the data distribution; black bars indicate the mean and gray bars indicate the median. Trials had to have a minimum of 1 pulse/s during baseline to be included in this dataset. Asterisks denote significant differences between driver and non-driver controls at each stimulation intensity (p < 0.05, Kruskal-Wallis test followed by a one-tailed Mann-Whitney U test, comparing driven to empty lines at the same intensity). Sample sizes through A-D: n = 16 flies of each genotype.

We next examined the wavelength dependence of song suppression, testing red, green, and blue light at comparatively low intensities to minimize behavioral effects of the light itself (Figure 5—figure supplement 2). Blue light was tested at 3.5–10 μW/mm², and green and red light at 2.1–6.2 μW/mm². Red light elicited robust inhibition of pulse song in flies expressing A1ACR1 or HfACR1, but not in those expressing GtACR1. In contrast, pulse song in empty-driver controls was only minimally affected (Figure 5—figure supplement 1A). Blue light (475 nm) caused modest suppression in flies expressing A1ACR1 and GtACR1, but also produced strong ACR-independent effects, rendering it unsuitable for studying singing behavior. Green light produced mild suppression in A1ACR1 and GtACR1, but at matched intensities, GtACR1 with green light was less effective than RubyACRs with red light. We also analyzed sine song production and found that red light slightly reduced sine song in A1ACR1 and HfACR1 flies, but not in GtACR1-expressing animals (Figure 5—figure supplement 3). These results indicate that RubyACRs are more effective than GtACR1 in suppressing neural activity during behavior.

Using red light in combination with RubyACRs to inhibit pIP10 activity offered several advantages over other wavelengths and tools such as GtACR1. The low-intensity light required for activation minimized unintended behavioral effects. Moreover, the inhibition was both robust and consistent across individuals within the treated populations. Activation of A1ACR1 led to a decrease in pulse production greater than 80% in 56% of flies at 6.2 μW/mm² red light, while HfACR1 resulted in a >80% reduction in 69% of animals. In contrast, neither GtACR1-expressing flies nor any control empty-driver flies had a decrease of more than 80%.

## Discussion

Our results demonstrate that RubyACRs are effective optogenetic inhibitors in *Drosophila melanogaster*. Electrophysiological recordings revealed strong hyperpolarization with rapid onset and offset kinetics in larval motor neurons. Functional imaging revealed decreases in calcium signals within the axonal arbor of cells in the visual system. Finally, three different behavioral assays demonstrated that these opsins are effective inhibitors in intact animals. These behavioral assays targeted a diverse set of neurons, from superficial glutamatergic motor neurons to neurons deep in the central brain: PPL1-γ1pedc and pIP10.

GtACR1 has been the standard optogenetic inhibitor in *Drosophila melanogaster* and has proven effective in many experimental protocols (Mauss et al., 2017, Mohammad et al., 2017). The GtACR1 action spectrum peaks in the green wavelengths (∼515 nm (Mohammad et al., 2017)), and accordingly, when we used red illumination (either 629 or 660 nm), we did not observe any evidence of hyperpolarization or strong behavioral effects in flies expressing GtACR1. By contrast, the red light sensitivity of RubyACRs is doubly beneficial—minimizing the effect of light on behavior and providing greater penetration through the fly’s cuticle to the nervous system (Inagaki et al., 2014a). Electrophysiological recordings further revealed that both A1ACR1 and HfACR1 elicited stronger and faster hyperpolarization than GtACR1, with HfACR1 generally outperforming A1ACR1. Notably, even when illuminating close to the optimal wavelength for GtACR1 activation, RubyACRs elicited responses at lower light intensities, highlighting their superior efficacy as optogenetic inhibitors.

The inhibition of courtship song with RubyACRs was both robust and spectrally specific: only red light suppressed song, while blue and green light had minimal effect at the tested intensities. This specificity may result from the relatively low light levels used in this assay, combined with the fact that shorter wavelengths are more strongly attenuated by tissue, and the target pIP10 neurons are located deep within the brain. This aligns with the broader observation that behavioral assays in intact flies tend to exhibit greater spectral sensitivity than electrophysiology and functional imaging of the exposed brain. The spectral specificity observed in the courtship assay may also stem from the use of 629 nm light, which closely matches the RubyACRs’ peak activation (∼610 nm). Using low-intensity, spectrally well-matched red light may be more effective for minimizing behavioral artifacts than attempting to minimize them using longer wavelengths at necessarily higher intensities.

Activating ACRs in axon terminals can lead to neurotransmitter release in mammalian neurons (Mahn et al., 2016). This arises due to an inversion of the chloride gradient in the axon, so although ACR opening hyperpolarizes the soma, it depolarizes the synaptic terminals and leads to neurotransmitter release (Messier et al., 2018). Our experiments did not reveal any evidence for RubyACR-induced neurotransmitter release. In particular, activating the neurons we targeted in the learning and courtship singing assays produces the opposite behavioral effects from what we observed: activation of PPL1-γ1pedc leads to aversive memory formation (Aso and Rubin, 2016), whereas we observed a weak appetitive memory; activation of pIP10 triggers song production (Bath et al., 2014; Calhoun et al., 2019; Clemens et al., 2018; Coen et al., 2016; Ding et al., 2019; Inagaki et al., 2014a; Roemschied et al., 2023; Shiozaki et al., 2024, 2024; von Philipsborn et al., 2011)), whereas we observed inhibition. Additionally, GCaMP imaging from axonal projections of visual system neurons showed a clear reduction in calcium levels upon RubyACR activation. Across a range of experimental conditions—including non-spiking neurons in the visual system—our findings consistently support the conclusion that RubyACRs inhibit neuronal signaling in Drosophila.

This suggests the possibility that an inverted chloride gradient in axons is not a feature of Drosophila neurons. Unlike highly polarized mammalian neurons, fly neurons exhibit less segregation between input and output sites (Schlegel et al., 2024), which may be accompanied by a more uniform chloride gradient across compartments. Another potential contributing factor could be that our experimental approaches likely activated RubyACRs across the entire neuron— including soma, dendrites, and axons—resulting in net hyperpolarization that prevents synaptic release. Of course, it remains possible that RubyACRs could trigger synaptic release in other contexts in Drosophila, as the polarity and/or steepness of the axonal chloride gradient may vary by neuron type or brain region. For example, activating GtACR1 or GtACR2 in larval cardiac tissue appears to have a depolarizing effect, as indicated by increased heart rate (Stanley et al., 2019). Similarly, in type III cold nociceptors, chloride currents are known to produce depolarization (Himmel et al., 2023). Fortunately, RubyACRs can readily be used in combination with green calcium indicators such as GCaMP to verify their effect on membrane polarization.

Overall, there was little to distinguish the performance of A1ACR1 from HfACR1, which differ by only five amino acids in the rhodopsin domain (Govorunova et al., 2020). That said, HfACR1 generates a stronger hyperpolarizing current than A1ACR1 (Govorunova et al., 2020), a difference that was evident in our recordings from larval motor neurons and generally reflected in behavioral assays, where HfACR1 typically elicited stronger effects. These included a persistent reduction in mobility when RubyACRs were activated in glutamatergic neurons. With each successive trial, activation of HfACR1 led to a progressive decline in baseline mobility, an effect clearly stronger than that observed with A1ACR1. This prolonged behavioral suppression is unlikely to result from an intrinsic property of the opsin itself—membrane potential changes in larval motor neurons occurred on the timescale of tens of milliseconds, far shorter than the persistence we observed. Instead, it more likely reflects a circuit-level mechanism. For example, sustained hyperpolarization of key glutamatergic neurons could disrupt downstream network activity in a manner that outlasts the light stimulus. Supporting this interpretation, long-lasting behavioral effects have also been reported with WiChR, a green light-activated potassium channel, in Drosophila (Ott et al., 2024). In contrast, when RubyACRs were expressed in pIP10 neurons, courtship song recovered rapidly and completely after stimulation, demonstrating that RubyACR-mediated inhibition can be transient depending on the circuit engaged.

The differences between A1ACR1 and HfACR1 provide useful options for circuit-level studies. A1ACR1 is likely to be effective in most applications, but HfACR1 may be preferable in more challenging contexts, such as deep tissue targets, low-expression regions, or neurons that are relatively resistant to hyperpolarization. As demonstrated, these effectors can be combined with sensors such as GCaMP to confirm whether hyperpolarization is occurring and to assess circuit-level consequences.

## Methods

### Transgenic animal construction

The coding sequences for the anion channelrhodopsins A1ACR1-EYFP and HfACR1-EYFP (Addgene #161025 and #161026) were codon-optimized for *Drosophila melanogaster* using GeneDesigner (ATUM). Synthesized with overlapping ends, the inserts were cloned into the plasmid backbones *pJFRC59-13XLexAop2-IVS-myr::GFP-p10* and *pJFRC82-20XUAS-IVS-Syn21-GFP-p10* (Pfeiffer et al., 2012) between the *NotI* and *XbaI* sites using Gibson assembly. Plasmid injections were carried out by GenetiVision Corporation.

### Fly Husbandry

Flies were reared on standard cornmeal molasses media supplemented with 0.2 mM all-trans retinal. At 2 to 3 days after eclosion, flies were selected and transferred to cornmeal molasses media supplemented with 0.4 mM all-trans retinal. Unless otherwise stated flies were kept at 25°C 40% humidity in dark only conditions.

### Measuring light intensity

Light intensity was measured using a Thorlabs SC170C microscope slide power sensor. For full-field illumination (e.g., in the Fly Bowl assay), the entire sensor area was used. Under the microscope objective, where light is confined to a small region, measurements were taken through a 200 μm diameter aperture cut into an aluminum sheet.

### Immunohistochemical Staining

Optogenetic actuators were crossed to RRa-GAL4, which also drove expression of tdTomato (pJFRC22-10XUAS-IVS-myr::tdTomato inserted in su(Hw)attP8). Third instar larvae were filleted. Following dissection, tissue was fixed using standard protocols (Aso et al., 2014). EYFP-labeled variants were detected with primary chicken anti-GFP (Invitrogen A10262, 1:1000) and secondary goat anti-chicken Alexa Fluor 488 Plus (Thermo Fisher A32931, 1:800). tdTomato was detected with primary rabbit anti-RFP (Clontech 632496, 1:1000) and secondary goat anti-rabbit Cy3 (Jackson 111-165-144, 1:1000).

### Electrophysiology

Optogenetic actuators were crossed into an RRa-GAL4 driver line that also carried GFP for visualizing aCC/RP2 motor neurons (*10XUAS-IVS-Syn21-GFP-p10* in *attP18::RRa-GAL4 / TM6B*). Third instar female larvae were dissected in ice-cold Schneider’s Insect Medium using a 35 mm Sylgard-coated Petri dish. The ventral nerve cord (VNC) was isolated, washed, and secured on a small piece of polylysine-coated coverslip. The coverslip was then transferred to a 1 mL perfusion chamber mounted under an upright microscope (Olympus BX61WI).

The preparation was continuously perfused at 0.5 mL/min with imaging buffer containing (in mM): NaCl 135, KCl 5, CaCl₂ 2, MgCl₂ 4, TES 5, sucrose 36; pH adjusted to 7.15 with NaOH. EYFP fluorescence was excited using a mercury light source (X-Cite exacte, Olympus) to locate RubyACR-expressing neurons.

A glass pipette (3–5 μm tip diameter) filled with 2% Type XIV protease (P5147, Sigma) in imaging buffer was positioned near the targeted cell. Pulses of positive pressure were used to enzymatically soften the glial sheath surrounding the soma. This was followed by alternating positive and negative pressure to remove debris and expose the neuron for recording.

Patch pipettes (resistance 5–8 MΩ) were pulled from borosilicate glass and filled with internal solution containing (in mM): K-gluconate 140, KCl 5, EGTA 2, MgCl₂ 2, HEPES 20; pH adjusted to 7.4 with KOH. Patch-clamp recordings were performed under transmitted light. Once whole-cell configuration was achieved, neurons were held at their resting membrane potential (*I* = 0).

Light pulses (490, 530, and 660 nm; durations from 1 msec to 30 s) were delivered with step-wise intensity increases via LEDs (Thorlabs part number, range tested μW/mm^2^):

- Blue (M490L4) : 7.8 to 300
- Green (M530L4) : 11 to 300
- Red (M660L4) : 3.7 to 274

Membrane potential was amplified using a MultiClamp 700B amplifier (Molecular Devices), digitized at 20 kHz via a USB-6343 I/O device (National Instruments), and recorded using WaveSurfer software (Taylor et al., 2025).

### Simultaneous 2-photon imaging and RubyACR activation

Imaging was performed on female flies using a resonant-scanning two-photon microscope (Spectra-Physics Insight DS DUAL) with near-infrared excitation at 920 nm and a 25× water-immersion objective (Nikon MRD77225 25XW). The system was controlled via ScanImage 2015.v3 (Vidrio Technologies). Images were acquired at a resolution of 512 × 512 pixels with a field of view of 141 × 141 μm and an approximate frame rate of ∼9 Hz (after averaging 5 frames). Excitation power at the sample was set to 12 mW for calcium imaging.

Photostimulation was delivered using a 660-nm LED (M660L3, Thorlabs) coupled to a digital micromirror device (DLPC300 LightCrafter, Texas Instruments) and introduced into the imaging path via a FF757-DiO1 dichroic mirror (Semrock). On the emission side, optical filtering included a primary dichroic (Di02-R635, Semrock), a detection path dichroic (565DCXR, Chroma), and emission filters FF03-525/50 and FF01-625/90 (Semrock).

Each trial consisted of a 10-second baseline period, a 20-second photostimulation phase, and a 20-second post-stimulation period. A total of 9 trials were conducted: trials 1–8 used increasing light intensities ranging from 35 to 1100 μW/mm², and trial 9 used a lower intensity of 88 μW/mm². Photostimulation intensity was measured using a Thorlabs S170C power sensor through a 200 μm aperture in an aluminum sheet, as described above.

Regions of interest (ROIs) were drawn over layers M8–M10 based on mean intensity projections. Fluorescence signals were processed using custom Python scripts. To calculate ΔF/F₀, the fluorescence offset was first subtracted, followed by subtraction of the baseline fluorescence (F₀), defined as the median fluorescence during the 5-second period preceding stimulation onset. The resulting ΔF was then normalized by F₀. Final traces were smoothed using a Gaussian filter (σ = 1).

### Walking behavior

Female flies were tested in the Fly Bowl behavioral arena as previously described (Branson et al., 2009; Robie et al., 2024, 2017). Two different stimulation protocols were used.

In the first protocol, each trial consisted of a 30-second baseline period followed by a 30-second stimulation period, during which flies were illuminated from below with a specified wavelength. Throughout both periods, 850 nm infrared illumination was maintained at 38 μW/mm² for video acquisition. Each protocol included 8 trials, with each intensity level tested in two consecutive trials before being increased.

For 660 nm stimulation, two intensity series were used:

- Series 1: 34% (60 μW/mm²), 50% (89 μW/mm²), 65% (120 μW/mm²), 80% (140 μW/mm²)
- Series 2: 6% (10 μW/mm²), 17% (29 μW/mm²), 34% (60 μW/mm²), 50% (89 μW/mm²) Data from both 660 nm protocols were pooled for analysis and figure presentation.

For other wavelengths, intensity steps were as follows:

- 475 nm: 40% (92 μW/mm²), 60% (130 μW/mm²), 80% (170 μW/mm²), 100% (190 μW/mm²)
- 525 nm: 40% (48 μW/mm²), 60% (68 μW/mm²), 80% (88 μW/mm²), 100% (110 μW/mm²)

The second protocol used a fixed 660 nm intensity of 34% (60 μW/mm²) with 10-second baseline and 10-second stimulation periods. This protocol included 21 trials.

### Olfactory arena

Flies were reared on food supplemented with all-trans-retinal (0.2 mM prior to eclosion, then 0.4 mM thereafter). Adult females aged 1–4 days were collected under light CO₂ anesthesia and sorted at least 24 hours prior to behavioral testing. Flies were not subjected to starvation.

Olfactory arena experiments were conducted as described previously (Aso and Rubin, 2016), with modifications to the odor delivery system: a two-step air-dilution olfactometer (Honegger et al., 2011)was used to supply odors. The final dilution for both 6-methyl-5-hepten-2-one and hexanal was 1:100 (v/v).

Behavior was recorded at 30 frames per second and tracked using Caltech FlyTracker (Eyjolfsdottir et al., 2014). Data were analyzed using custom MATLAB scripts.

### Recording courtship song

Adult males were isolated within 6 hours of eclosion and housed individually for 4–7 days before testing. Virgin females were aged 3-10 hours post-eclosion. Experiments were conducted using Song Torrent, a high-throughput audio and video recording system (Sawtelle et al., 2024). A single male and a single female were placed together in a recording chamber. RubyACR was expressed in pIP10 neurons using the driver line VT040556-p65ADZp@attp40; VT040347-ZpGDBD@attp2.

Two experimental protocols were used to assess courtship behavior. Each trial consisted of a 5-second baseline, a 5-second testing period, and a 5-second post-stimulation period. During the testing period, the chamber was continuously illuminated with an LED emitting a specific wavelength, while ambient illumination from a white LED was maintained throughout the experiment.

In the first protocol, illumination from a 629 LED was tested over six different intensities: 4.2, 8.3, 17, 25, 33, and 41 μW/mm^2^. Intensities were tested step wise, from lowest to highest and then this stepwise protocol was repeated 12 times.

In the second protocol, three different wavelengths (475, 530, and 629 nm) were tested at three different intensities. Again, intensities were tested stepwise, with the lowest intensity for each wavelength tested first, and then the cycle was repeated 8 times. Illumination intensities were as follows:

- 475 nm: 3.5, 7, and 10 μW/mm²
- 530 nm and 629 nm: 2.1, 4.2, and 6.2 μW/mm²

Courtship song events were detected in the audio recordings using SongExplorer, a machine learning-based platform for segmenting acoustic communication signals (Arthur et al., 2021). *D. melanogaster* produce two distinct song types during courtship: sine and pulse songs (Schilcher, 1976). Using recordings from wild-type flies without optogenetic stimulation, we manually annotated pulse song, sine song, inter-pulse intervals, non-song noise, and ambient noise as described previously (Shiozaki et al., 2024). These annotations were then used to train a classification model.

## Data & Reagent Availability

The data and code used to produce the figures are deposited at https://doi.org/10.5061/dryad.gxd2547zg

Fly strains constructed during this work are available at the Bloomington Drosophila Stock Center, stock ID numbers: 606146-606153.

## Acknowledgements

We thank the GENIE Steering Committee, which at the time of this work consisted of: Jeremy Hasseman, Vivek Jayaraman, Wyatt Korff, Eric Schreiter, Nelson Spruston, Alison Tebo and Glenn Turner for their guidance and oversight. We thank David Stern for the use of his lab’s equipment for courtship song experiments. We thank Christina Christoforou (Janelia Project Technical Resources) for performing larval dissections, immunohistochemistry, and confocal imaging. We thank Vasily Goncharov and Janelia Experimental Technology (jET) for imaging support and resources. We thank Katie Schretter, Alice Robie and Adam Taylor for assistance with Fly Bowl experiments and analysis. We are grateful to Yoshinori Aso for his feedback on the manuscript. ChatGPT, Claude, and Perplexity AI services were used to suggest coding examples and writing revisions for this work.

---

**Figure 1—figure supplement 1:**
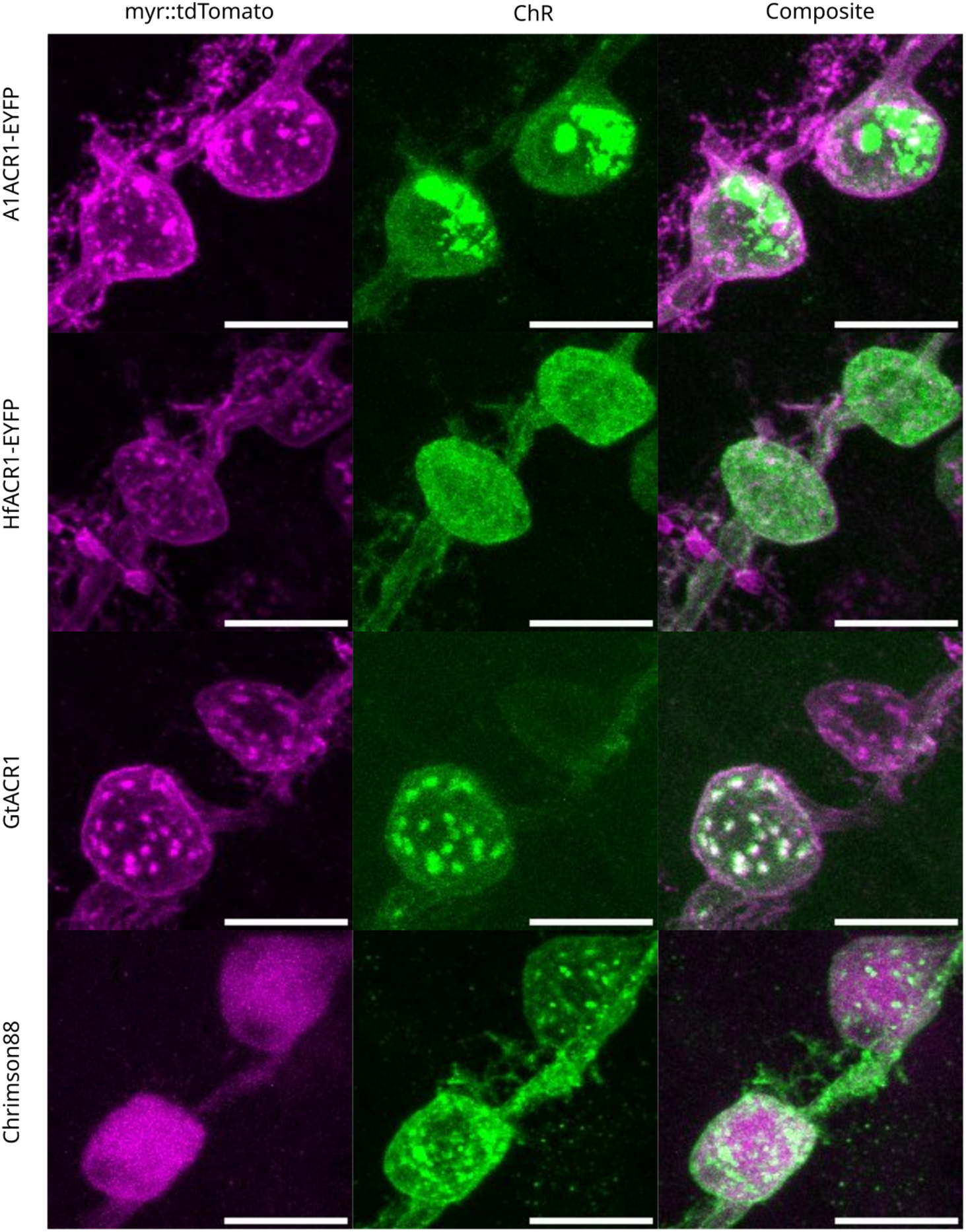
RubyACR-EYFP expression in larval motor neuron cell bodies. Green shows staining for the ACR variants and Chrimson, magenta is the membrane localized myr::tdTomato and white represents colocalization; all images are maximum intensity projections (scale bar = 10 μm). Rows correspond to different variants, driver is RRa-GAL4. Puncta are visible in all cases except HfACR1-EYFP.

**Figure 3—figure supplement 1:**
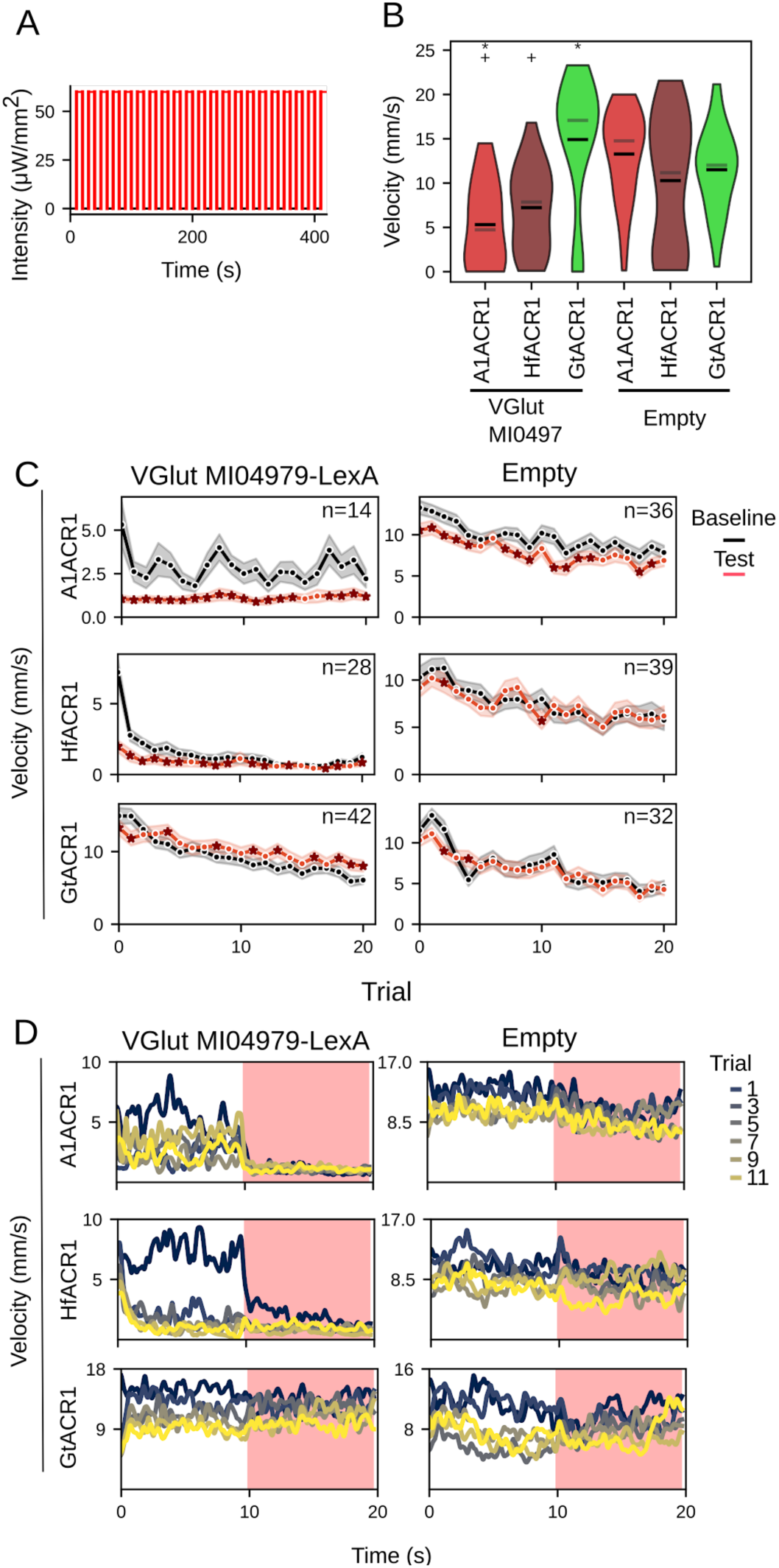
Optogenetically inhibiting spontaneous locomotor activity with LexAop2-RubyACR variants. A. Protocol spans 21 trials with each trial having baseline (black) and testing (red) periods when the chamber is illuminated at 29 µW/mm^2^ with a 660 nm LED. Both baseline and testing period occurs over 10 sec periods. OR Stimulation protocol: each trial consists of a 10-second baseline period (black) followed by a 10-second testing period (red), during which the chamber is illuminated with 660 nm light at 29 μW/mm². This cycle is repeated for 21 trials. B. Walking velocity during the baseline period of the first trial, prior to any LED stimulation. Asterisks indicate driver–effector genotypes with significantly lower velocity than their empty-driver controls (p < 0.05, Kruskal-Wallis test with post hoc Mann-Whitney U test). A plus symbol (+) denotes a significant difference between GtACR1 and other ChRs expressed with the same driver (p < 0.05). C. Mean velocity in each trial. Black lines show average velocity during baseline; red lines show velocity during the subsequent illuminated testing period. As indicated, rows correspond to a specific ChR and columns to a driver. Asterisks as markers denote trials with a significant difference between baseline and stimulation periods (p < 0.05, Kruskal-Wallis test with post hoc Wilcoxon signed-rank test). Numbers of flies tested are indicated in the right top corner of each plot. D. Velocity measured per second during trials 1, 3, 5, 7, 9, and 11, as indicated in the legend. Red shaded region indicates the illumination during the testing period. ChR variants are arranged along the rows, and drivers along the columns.

**Figure 5—figure supplement 1:**
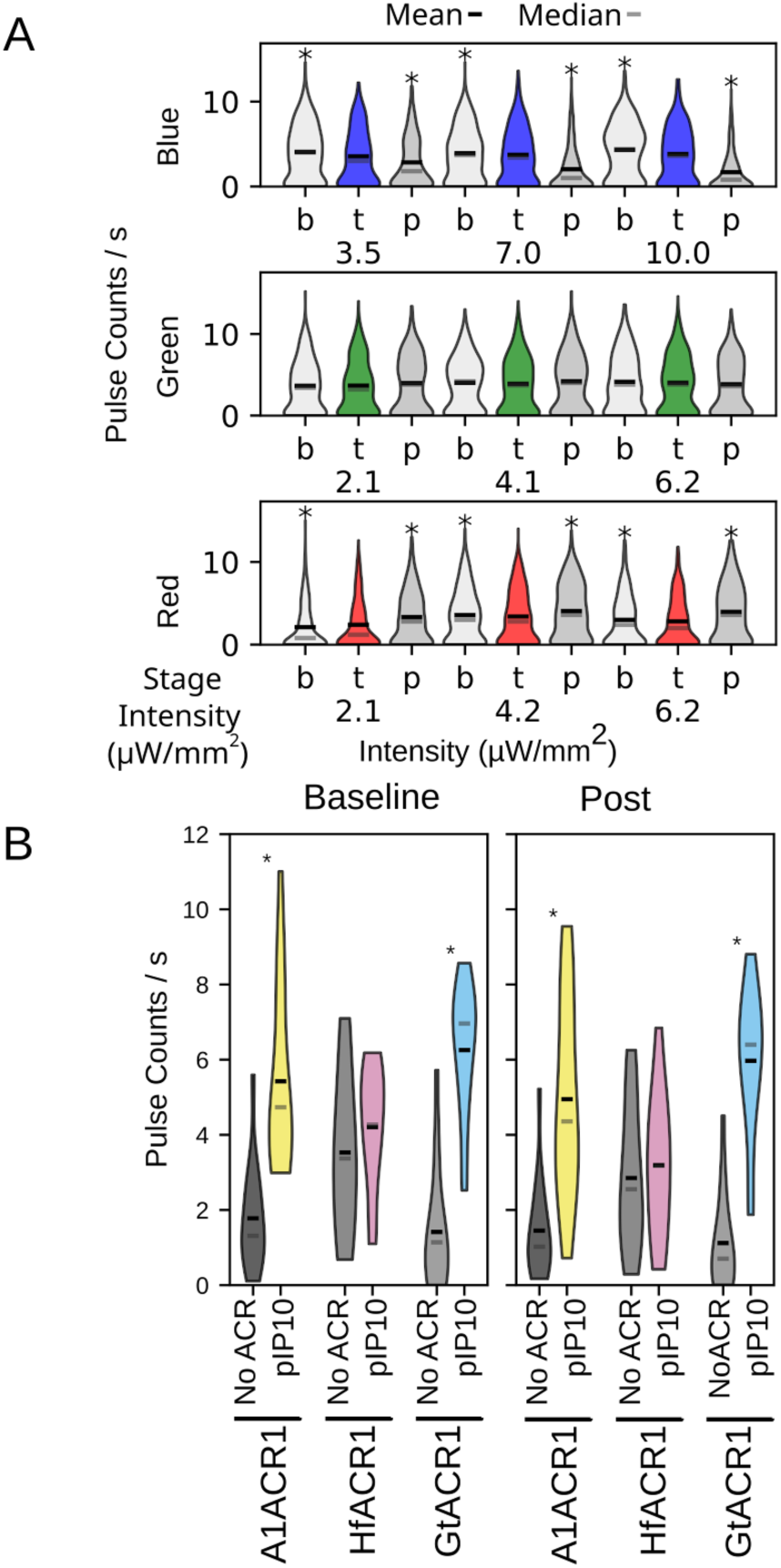
The effects of light and opsin expression on pulse song. A. Mean pulse counts per second of control, empty-driver flies (with or without the ChR transgene) for each trial stage across different stimulation wavelengths and intensities. Stages are: baseline (b) prior to illumination; testing (t) when illuminated and post-testing (p). Stimulation LEDs had peak wavelengths at 475 nm (blue, top row), 530 nm (green, middle row), and 629 nm (red, bottom row). Both blue and red light elicited significant changes in pulse frequency in these control flies (p < 0.05, Kruskal-Wallis test with post-hoc Wilcoxon signed-rank test; n = 64 flies). B. A1ACR1 and GtACR1 expression increases pulse song. Pulse counts are compared between flies with and without the pIP10 driver (n = 16 per condition). For both baseline and post-stimulation periods, data are pooled across all tested wavelengths and intensities for each genotype. Gray bars represent flies lacking the pIP10 driver; colored bars represent flies with ChR expression driven by pIP10. Asterisks indicate significant differences between empty-driver and pIP10-driver flies (p < 0.05, Kruskal-Wallis test with post-hoc Wilcoxon signed-rank test).

**Figure 5—figure supplement 2:**
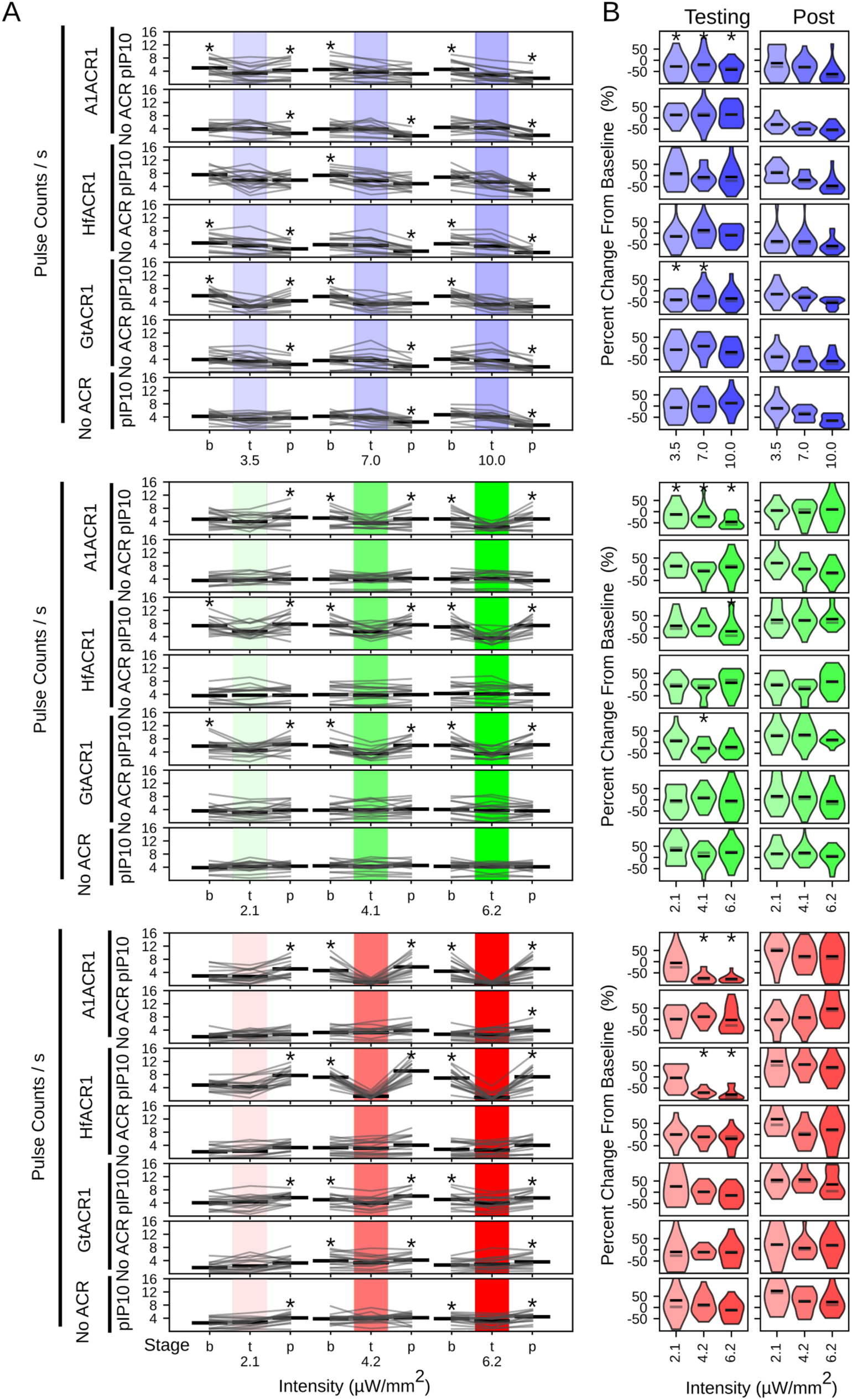
Effects of different stimulation wavelengths and light intensities on pulse song. A. Mean pulse counts per second for each trial stage for different stimulation wavelengths and intensities. LEDs with peak illumination at 475 (blue, top), 530 (green, middle), and 629 nm (red, bottom) were tested. Each row represents a particular driver-effector combination, as indicated at left. Stages are 5-second epochs of: baseline (b) prior to illumination; testing (t) when illuminated and post-testing (p). Illumination intensity increased with each successive trial as indicated by the shading. Gray lines represent data from individual courting pairs (n = 16), black bars indicate mean over all pairs. Asterisks denote significant differences between the testing period and either the baseline or post period (p < 0.05, Kruskal-Wallis test with post-hoc Wilcoxon signed-rank test). B. Percent change in pulse song frequency from baseline to testing (left) and from baseline to post (right). Driver-effector combinations for each row continue from **A**. Violin plots reflect the data distribution; black bars indicate the mean and gray bars indicate the median. Trials had to have a minimum of 1 pulse/s during baseline to be included in this dataset. Asterisks denote significant differences between driver and non-driver controls at each stimulation intensity (p < 0.05, Kruskal-Wallis test with post-hoc Mann-Whitney U test). Sample sizes in A,B: n = 16 flies of each genotype.

**Figure 5—figure supplement 3:**
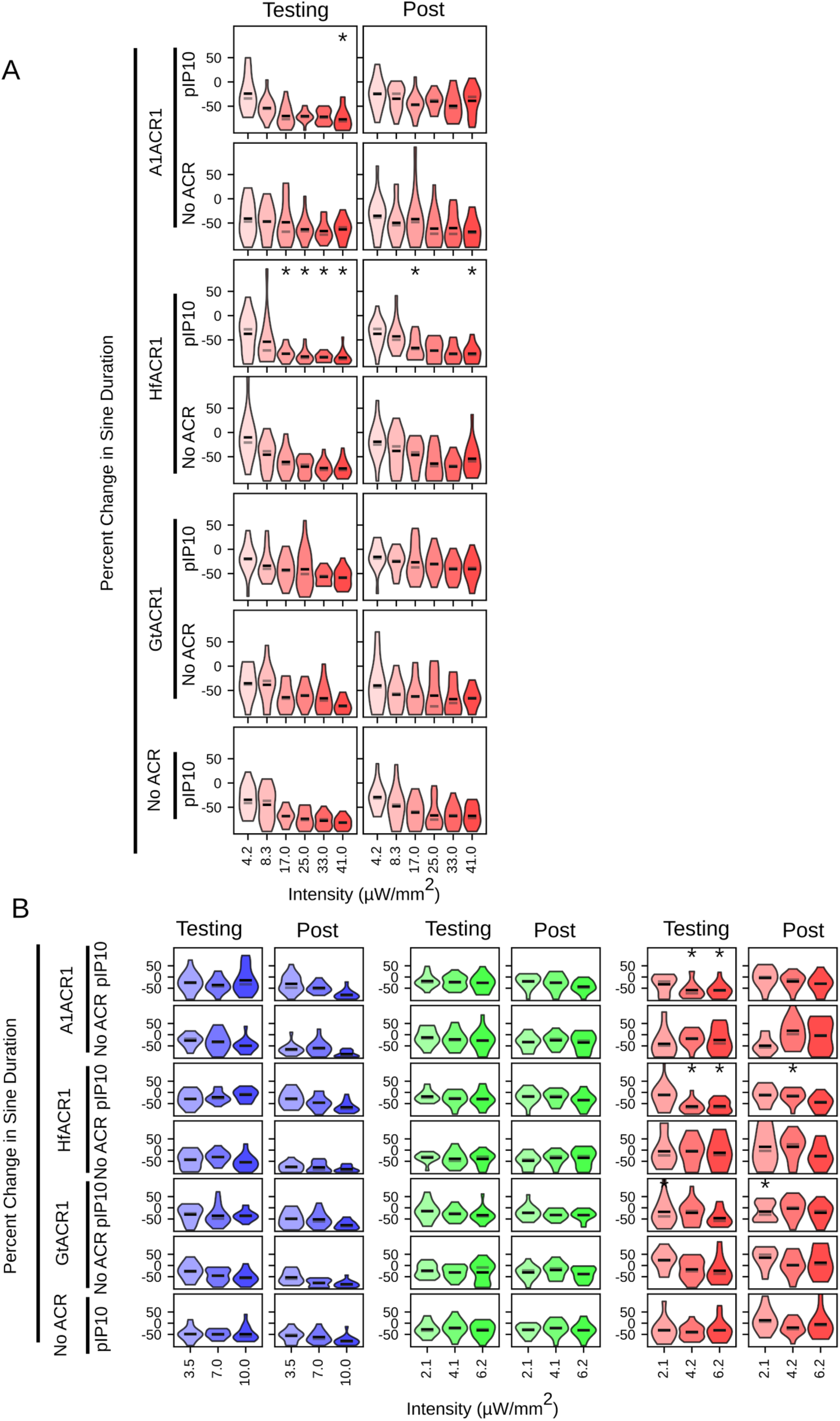
Activating RubyACRs in pIP10 does not consistently suppress sine song duration. A. Percent change in sine song duration from baseline to testing (left) and from baseline to post (right) when illuminated with a red (660 nm) LED. As indicated, each row represents a particular driver-effector combination. Red LED intensity increases from left to right. Violin plots reflect the data distribution; black bars indicate the mean and gray bars indicate the median. Asterisks denote significant differences between driver and non-driver controls at each stimulation intensity (p < 0.05, Kruskal-Wallis test with post-hoc Mann-Whitney U test). B. Percent change in sine song when illuminated with either blue (475 nm), green (530 nm), or red (660 nm) as indicated by background color. Intensities for each color increase from left to right. Percent change between baseline and testing is indicated on the left while percent change from baseline to post is indicated on the right. Asterisks denote significant differences between driver and non-driver controls at each stimulation intensity (p < 0.05, Kruskal-Wallis test with post-hoc Mann-Whitney U test). Sample sizes in A,B: n = 16 flies of each genotype.

## Abbreviations used in text

msec: millisecond
μW/mm²: microWatt per millimeter squared

## Transgenes used in experiments

| Transgene | Source | Notes | Figure |
| --- | --- | --- | --- |
| pJFRC22-10XUAS-IVS-myr::tdTomato in su(Hw)attP8 | (Pfeiffer et al., 2010) |  | 1 |
| pGP-20XUAS-IVS-Syn21-op1-A1ACR1_EYFP-p10.627.1 in attP18 |  |  | 1,2,3,4,5 |
| pGP-20XUAS-IVS-Syn21-op1-HfACR1_EYFP-p10.628.1 in attP18 |  |  | 1,2,3,4,5 |
| 20XUAS-opGtACR1-kir-mVenus in attP18 |  |  | 1,3,5 |
| 10XUAS-IVS-Syn21-Chrimson88-tdT-3.1 in attP18 | (Klapeetke et al., 2014) |  | 1,2 |
| pJFRC81-10XUAS-IVS-Syn21-GFP-p10 in attP18 | (Pfeiffer et al., 2012) |  | 1 |
| 19F01-Gal4 (attP2) | (Dionne et al., 2018) | Drives expression in Mi1 neurons | 2 |
| pGP-20XUAS-IVS-Syn21-59D.1Ldr-CsChrimson-KIR2.1-mVenus-p10.612.1 in attP18 | (Klapeetke et al., 2014) |  | 2 |
| 20XUAS-Syn21-opGCaMP6s in su(Hw)attP8 | (Chen et al., 2013) |  | 2 |
| VGlut MI04979-Gal4 | (Diao et al., 2015) |  | 3 |
| VGLUT-Gal4 | (Deng et al., 2019) |  | 3 |
| VGlut MI04979 LexA:QFAD | (Diao et al., 2015) |  | 3 |
| MB320C/MB320C/w;; TH-p65ADZp in VK00027, 22B12-ZpGdbd in attP2/TM3/w[1118]; P{y[+t7.7]w[+mC]=R22B12-GAL4.DBD}attP2 PBac{y[+mDint2]w[+mC]=ple-p65.AD}VK00027 | (Hige et al., 2015) |  | 4 |
| VT040556-p65ADZp@attP40; VT040347-ZpGDBD@attP2 | (Ding et al., 2019) | Drives expression in pIP10 | 5 |

